## Supplementary material 4 for "Creating simple predictive models in ecology, conservation and environmental policy based on Bayesian belief networks"

### Running BBNet Functions

2024-02-28

### BBNet

#### Installing the BBNet package

The BBNet package is available from CRAN and can be installed (if required) and loaded using the following code

```
if (!require("bbnet")) install.packages("bbnet")
```

```
## Loading required package: bbnet

## Loading required package: dplyr

##
## Attaching package: 'dplyr'

## The following objects are masked from 'package:stats':
##
##     filter, lag

## The following objects are masked from 'package:base':
##
##     intersect, setdiff, setequal, union

## Loading required package: ggplot2

## Loading required package: grid

## Loading required package: igraph

##
## Attaching package: 'igraph'

## The following objects are masked from 'package:dplyr':
##
##     as_data_frame, groups, union

## The following objects are masked from 'package:stats':
##
##     decompose, spectrum
```

```
## The following object is masked from 'package:base':
##
##     union
```

```
## Loading required package: tibble
```

```
##
## Attaching package: 'tibble'
```

```
## The following object is masked from 'package:igraph':
##
##     as_data_frame
```

#### Importing data

With the exception of the bbn.network.diagram() function, all functions require a network model in the format below. It is easiest to read these into the R environment from a csv file. Initially we will open the Rocky Shore model

```
setwd("G:\\My Drive\\MainDocs\\Writing in progress\\BBN final stuff\\R Code and Data\\SuppMat\\SuppMat1

my_BBN <- read.csv('RockyShoreNetwork.csv', header=T)

head(my_BBN)
```

```
##             X Dogwhelk Topshell Limpet Periwinkle Barnacle Green.Algae Biofilm
## 1    Dogwhelk       NA       -1     -2         -2       -3          NA      NA
## 2    Topshell       NA       NA     -1         -1       NA          -3      -3
## 3      Limpet       NA       -2     NA         -2       NA          -4      -4
## 4  Periwinkle       NA       -1     -1         NA       NA          -3      -3
## 5    Barnacle       NA       NA     NA         NA       NA          NA      NA
## 6 Green Algae       NA       NA     NA         NA       NA          NA      -2
##   Corline.algae Fucoid.Algae
## 1            NA           NA
## 2            NA           NA
## 3            NA           NA
## 4            NA           NA
## 5            NA           NA
## 6            -3           -1
```

The details of this interaction matrix are discussed in the main text of the paper. Essentially it explains direct interactions between what is here the species or taxon in the *row* on the species or taxon in the *column.*

It is also normal in most functions to have a scenario or scenarios which we want to investigate. In this rocky shore example, we have three scenarios - the removal of dogwhelks, the addition of periwinkles and a combination treatment, where dogwhelks are removed and periwinkles added. Some of these scenarios and the model are described in more detail in this paper.

Scenarios are loaded in as follows:

```
dogwhelk <- read.csv('Dogwhelk_Removal.csv', header = T)
winkle <- read.csv('Winkle_addition.csv', header = T)
combined <- read.csv('Combined_Treatment.csv', header = T)

head(dogwhelk)
```

```
##   Increase        Node
## 1       -4    Dogwhelk
## 2        0    Topshell
## 3        0      Limpet
## 4        0  Periwinkle
## 5        0    Barnacle
## 6        0 Green Algae
```

In this case, any planned manipulation of the system (i.e. direct effects on the system) are listed - here all dogwhelks are removed, so a value of -4 is given (scale from -4 -strong decrease, to +4, strong increase).

#### Running a predictive model

#### bbn.predict()

We can run a predictive model on up to 12 scenarios at a time using the bbn.predict() function. As a minumum, we need to pass the network model and one scenario to the function.

```
bbn.predict(bbn.model = my_BBN, priors1 = dogwhelk, figure = 0) # figure set to zero, this is explained
```

```
## [1] "Scenario number  1"
##     Increase          name    LowerCI    UpperCI
## 1 -4.0000000      Dogwhelk -4.0000000 -4.0000000
## 2  0.8000000      Topshell  0.8000000  0.8000000
## 3  1.6000000        Limpet  1.6000000  1.6000000
## 4  1.6000000    Periwinkle  1.6000000  1.6000000
## 5  2.4000000      Barnacle  2.4000000  2.4000000
## 6 -1.7985382   Green Algae -1.7985382 -1.7985382
## 7 -1.7985382       Biofilm -1.7985382 -1.7985382
## 8  1.0791229 Corline algae  1.0791229  1.0791229
## 9  0.3597076  Fucoid Algae  0.3597076  0.3597076
```

The output given above shows the *posterior* outcome of the model as a result of dogwhelks decreasing - under the Increase column. Grazers increase (positive values) and some of the seaweeds which would be grazed by the grazers decreases. As the paper above explains, this is a short-term prediction model, so some seaweed actually increases in this scenario. There are also confidence intervals of our predictions - currently these show no difference to the main values, but we can bootstrap the model to gain an idea of the confidence of the predictions.

###### Function arguments

*Required arguments*

*bbn.model* - a matrix or dataframe of interactions between different model *nodes* *priors1* - an X by 2 array of initial changes to the system under investigation. The first column should be a -4 to 4 (including 0) integer value for each node in the network with negative values indicating a decrease and positive values representing an increase. 0 represents no change. Note, names included here are included as outputs in tables and figures (note misspelling of coraline algae in examples). Shortening these names can provide better figures

*Optional Arguments*

*priors2* - *priors12* - as above, but additional scenarios.

*boot_max* - the number of bootstraps to perform. Suggested range for exploratory analysis 1-1000. For final analysis recommended size = 1000 - 10000 - note, this can take a long time to run. Default value is 1, running with no bootstrapping - suitable for exploration of data and error checking.

*values* - default value 1. This provides a numeric output of posterior values and any confidence intervals. Set to 0 to hide this output

*figure* - default value 1. Sets the figure options. 0 = no figures produced. 1 = figure is saved in working directory as a PDF file (note, this is overwritten if the name is not changed, and no figure is produced if the existing PDF is open when the new one is generated). 2 = figure is produced in a graphics window. All figures are combined on a single plot where scenario 2 is below scenario 1 (i.e. scenarios work in columns then rows)

*font.size* - default = 5. This sets the font size on the figures.

*Example*

```
bbn.predict(bbn.model = my_BBN, priors1 = dogwhelk, priors2 = winkle, priors3= combined, figure = 2, boo
```

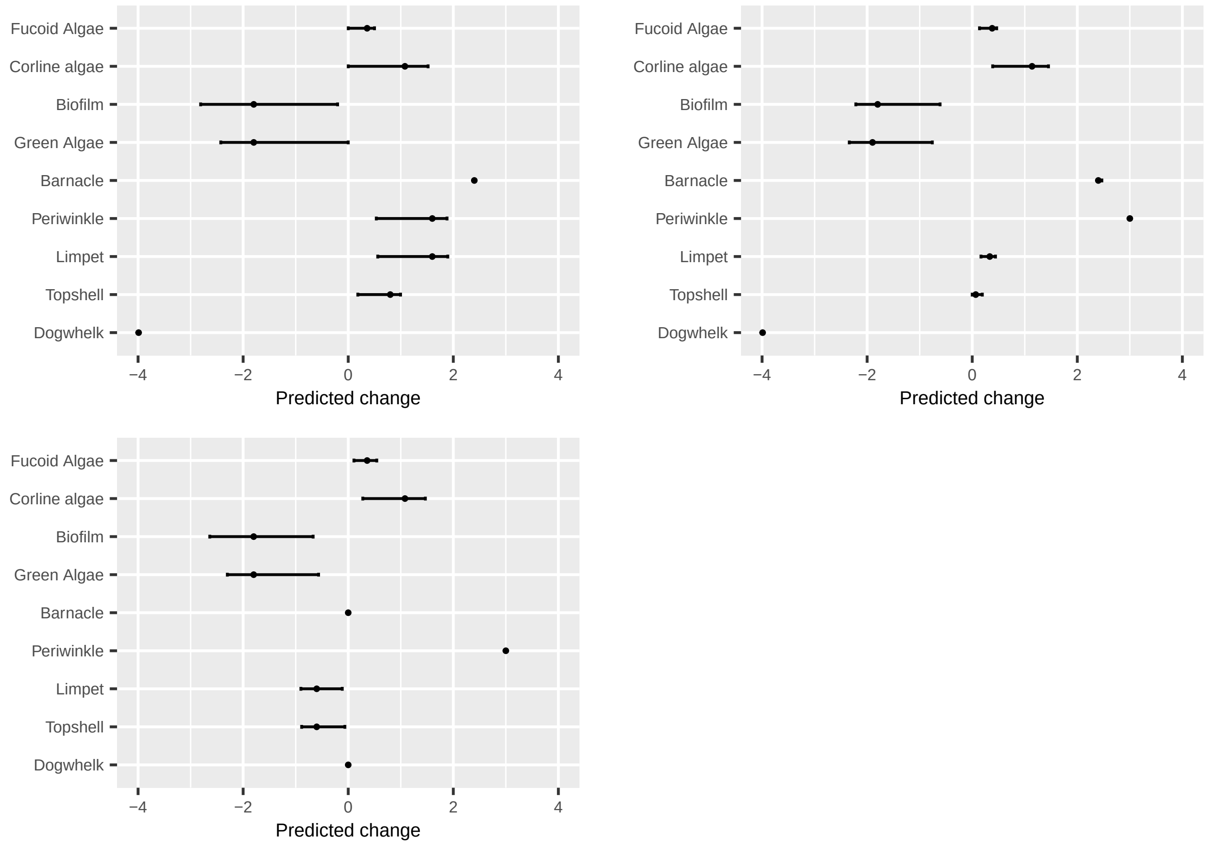

#### Visualising changes over time

Two functions help visualise changes over time. It should be noted that the exact values from these functions do not correspond to the more robust bbn.predict() - which should be used to inform of likely changes. These help visualise the flow of information through the network and how changes progress through the network over time (e.g. should we expect to see a change in one parameter before another - perhaps as per trophic cascade or ecological succession type processes)

#### bbn.timeseries()

As above we need to pass the function a network model and a scenario as a minimum. In this case, only one scenario can be analysed at once. The output is a graph of each node in the network, visualised across the different timesteps in the model. Note - values are plotted on each graph and lines of best fit are drawn using the geom_smooth() function. Typically this function may not perform well with the variability in values and lack of data points, and multiple warning messages may be produced.

*Required arguments*

*bbn.model* - a matrix or dataframe of interactions between different model *nodes*

*priors1* - an X by 2 array of initial changes to the system under investigation. The first column should be a -4 to 4 (including 0) integer value for each node in the network with negative values indicating a decrease and positive values representing an increase. 0 represents no change.

*Optional Arguments*

*timesteps* - default = 5. This is the number of timesteps the model performs. Note, timesteps are arbitrary and non-linear. However, something occurring in timestep 2, should occur before timestep 3.

*disturbance* - default = 1. 1 - creates a prolonged or press disturbance as per the bbn.predict() function. Essentially prior values for each manipulated node are at least maintained (if not increased through reinforcement in the model) over all timesteps. 2 - shows a brief pulse disturbance, which can be useful to visualise changes as peaks and troughs in increase and decrease of nodes can propagate through the network

*Example*

```
bbn.timeseries(bbn.model = my_BBN, priors1 = combined, timesteps = 5, disturbance = 2)
```

```
## ‘geom_smooth()‘ using formula = ’y ~ x’

## Warning in simpleLoess(y, x, w, span, degree = degree, parametric = parametric,
## : span too small.  fewer data values than degrees of freedom.

## Warning in simpleLoess(y, x, w, span, degree = degree, parametric = parametric,
## : pseudoinverse used at 0.98

## Warning in simpleLoess(y, x, w, span, degree = degree, parametric = parametric,
## : neighborhood radius 2.02

## Warning in simpleLoess(y, x, w, span, degree = degree, parametric = parametric,
## : reciprocal condition number 0

## Warning in simpleLoess(y, x, w, span, degree = degree, parametric = parametric,
## : There are other near singularities as well. 4.0804
```

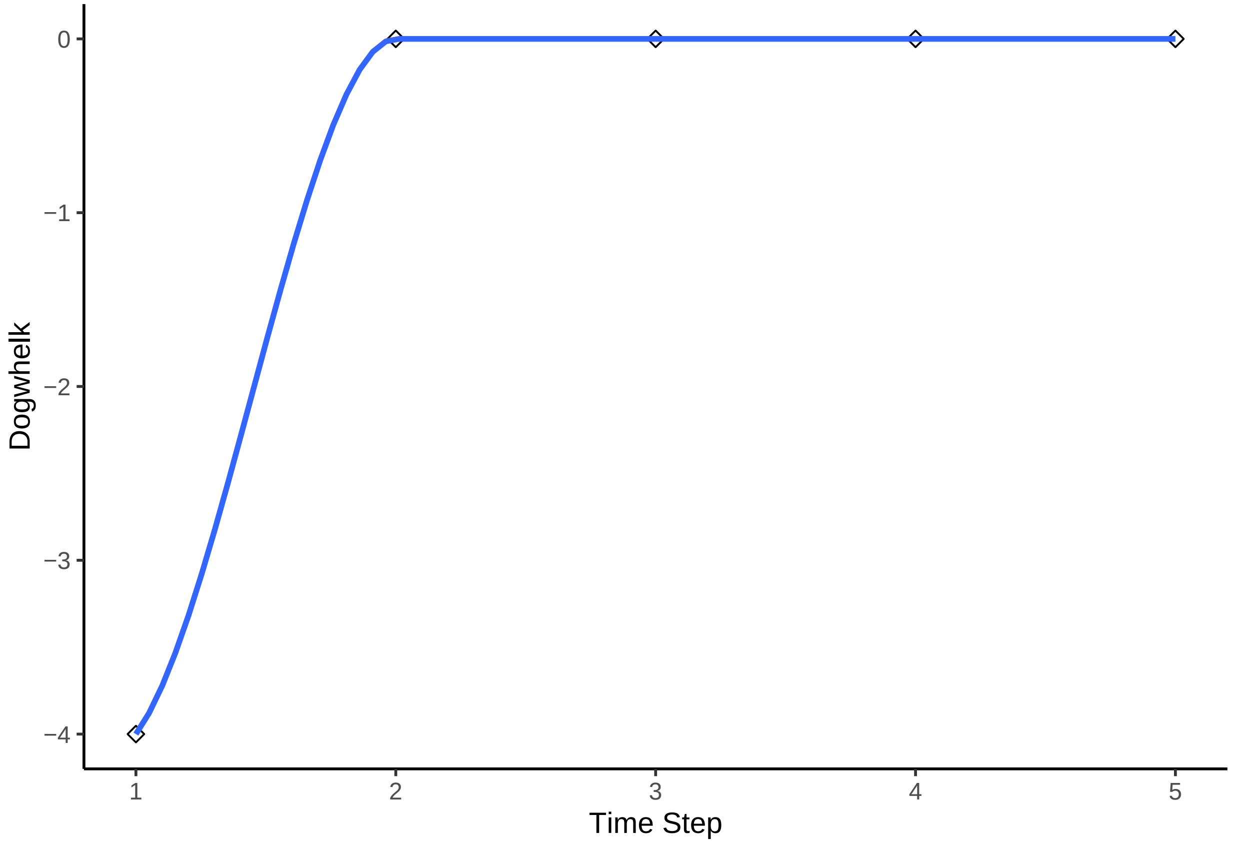


```
## 'geom_smooth()' using formula = 'y ~ x'

## Warning in simpleLoess(y, x, w, span, degree = degree, parametric = parametric,
## : span too small.  fewer data values than degrees of freedom.

## Warning in simpleLoess(y, x, w, span, degree = degree, parametric = parametric,
## : pseudoinverse used at 0.98

## Warning in simpleLoess(y, x, w, span, degree = degree, parametric = parametric,
## : neighborhood radius 2.02

## Warning in simpleLoess(y, x, w, span, degree = degree, parametric = parametric,
## : reciprocal condition number 0

## Warning in simpleLoess(y, x, w, span, degree = degree, parametric = parametric,
## : There are other near singularities as well. 4.0804
```

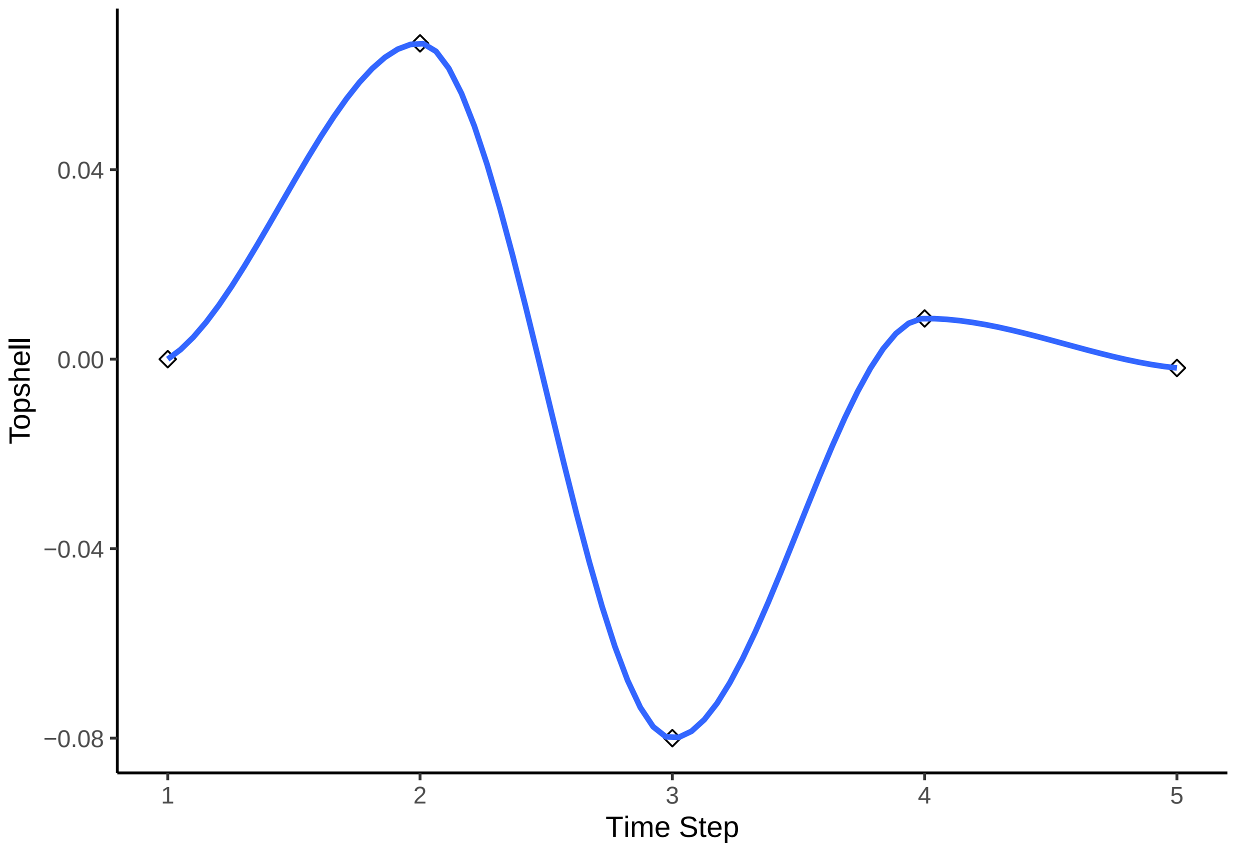


```
## 'geom_smooth()' using formula = 'y ~ x'

## Warning in simpleLoess(y, x, w, span, degree = degree, parametric = parametric,
## : span too small.  fewer data values than degrees of freedom.

## Warning in simpleLoess(y, x, w, span, degree = degree, parametric = parametric,
## : pseudoinverse used at 0.98

## Warning in simpleLoess(y, x, w, span, degree = degree, parametric = parametric,
## : neighborhood radius 2.02

## Warning in simpleLoess(y, x, w, span, degree = degree, parametric = parametric,
## : reciprocal condition number 0

## Warning in simpleLoess(y, x, w, span, degree = degree, parametric = parametric,
## : There are other near singularities as well. 4.0804
```

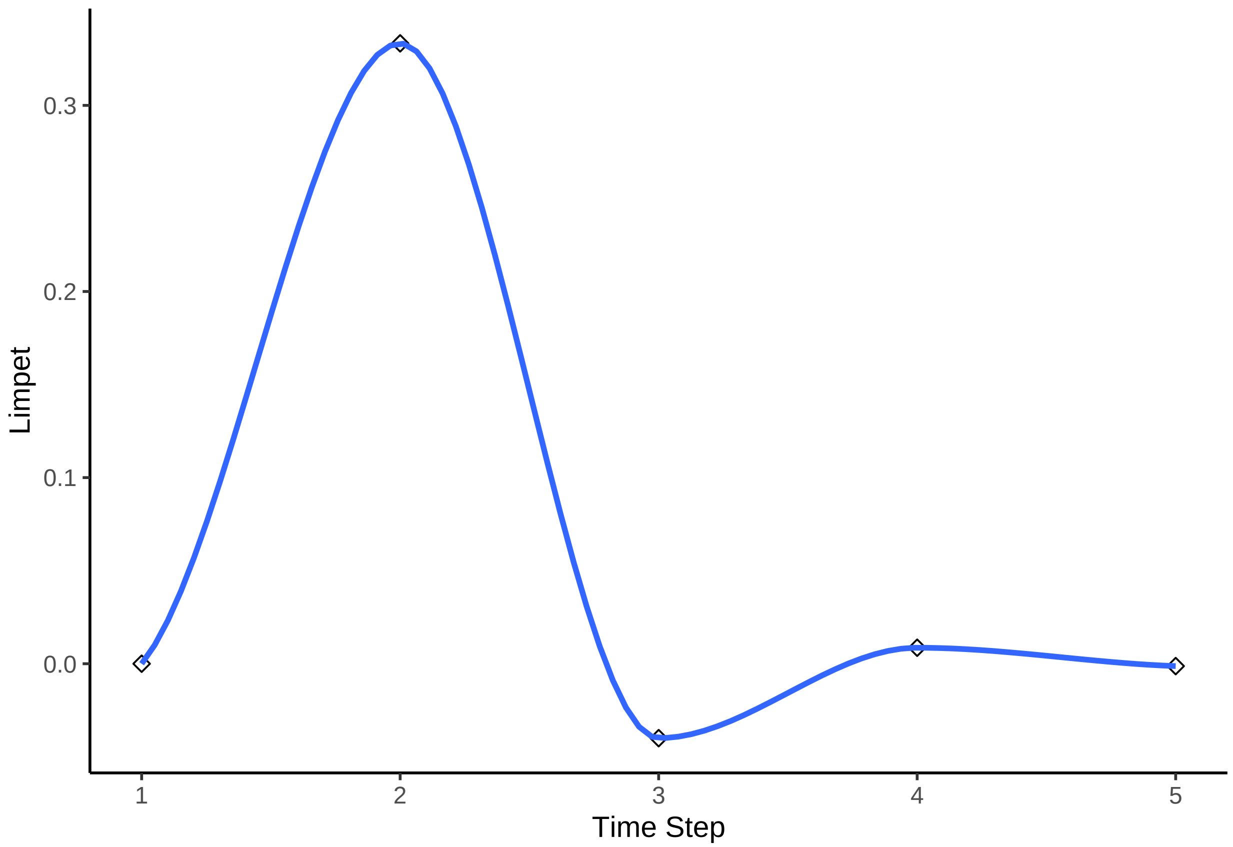


```
## `geom_smooth()` using formula = 'y ~ x'

## Warning in simpleLoess(y, x, w, span, degree = degree, parametric = parametric,
## : span too small.  fewer data values than degrees of freedom.

## Warning in simpleLoess(y, x, w, span, degree = degree, parametric = parametric,
## : pseudoinverse used at 0.98

## Warning in simpleLoess(y, x, w, span, degree = degree, parametric = parametric,
## : neighborhood radius 2.02

## Warning in simpleLoess(y, x, w, span, degree = degree, parametric = parametric,
## : reciprocal condition number 0

## Warning in simpleLoess(y, x, w, span, degree = degree, parametric = parametric,
## : There are other near singularities as well. 4.0804
```

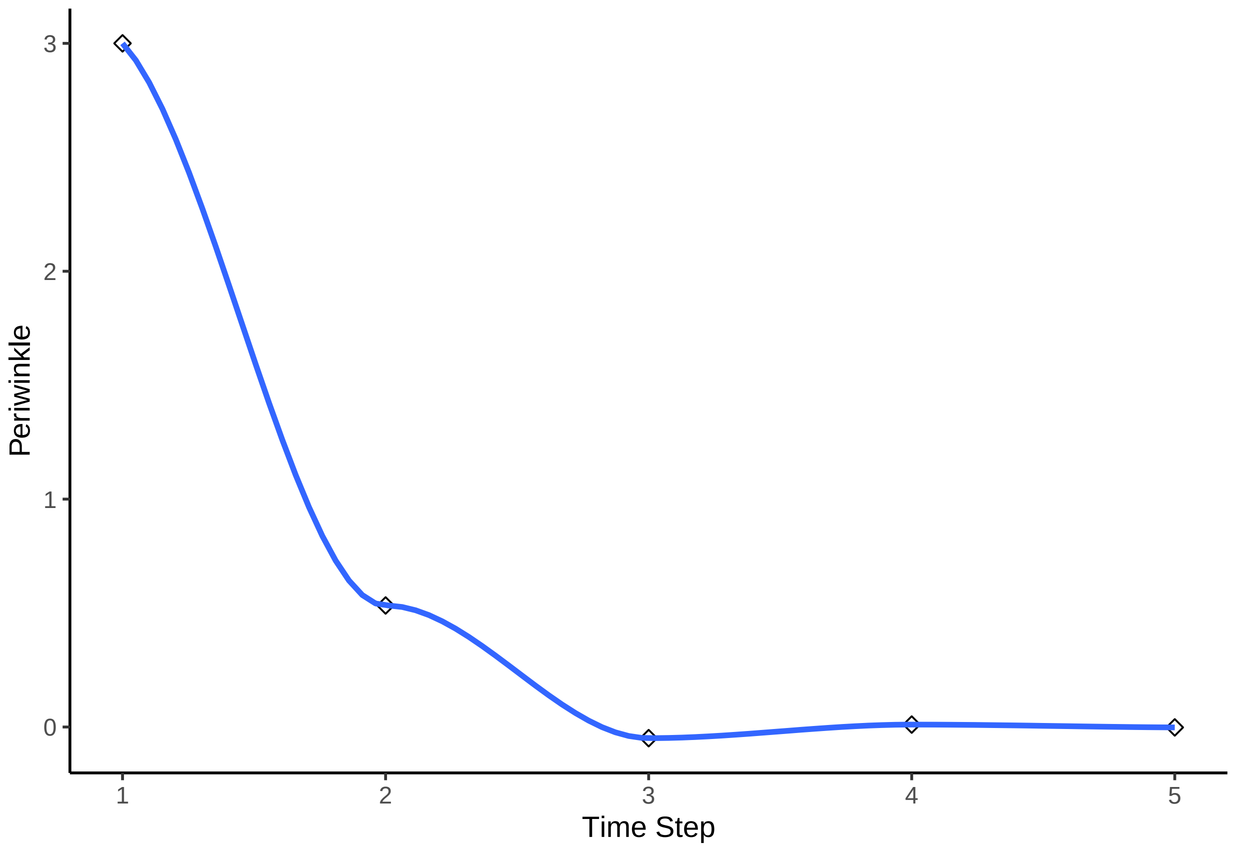


```
## `geom_smooth()` using formula = 'y ~ x'

## Warning in simpleLoess(y, x, w, span, degree = degree, parametric = parametric,
## : span too small.  fewer data values than degrees of freedom.

## Warning in simpleLoess(y, x, w, span, degree = degree, parametric = parametric,
## : pseudoinverse used at 0.98

## Warning in simpleLoess(y, x, w, span, degree = degree, parametric = parametric,
## : neighborhood radius 2.02

## Warning in simpleLoess(y, x, w, span, degree = degree, parametric = parametric,
## : reciprocal condition number 0

## Warning in simpleLoess(y, x, w, span, degree = degree, parametric = parametric,
## : There are other near singularities as well. 4.0804
```

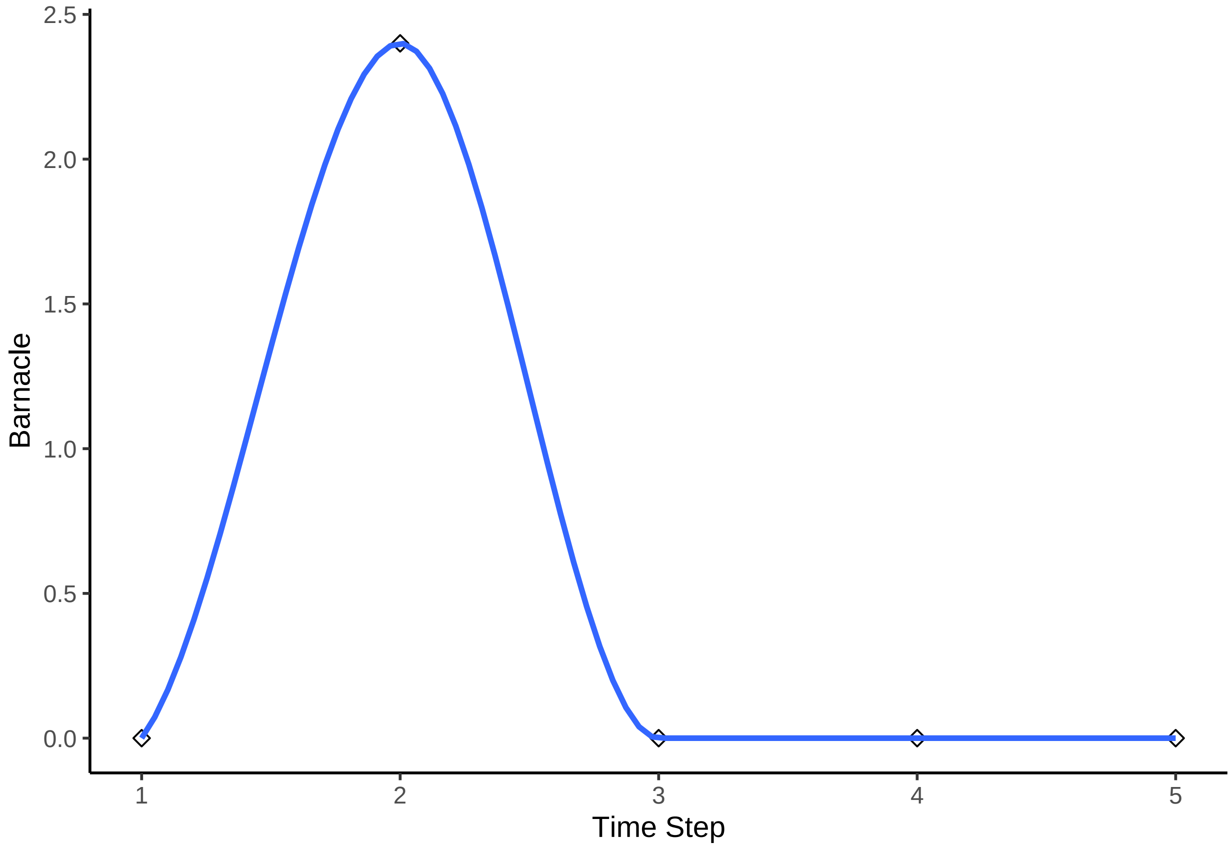


```
## ‘geom_smooth()‘ using formula = ’y ~ x’

## Warning in simpleLoess(y, x, w, span, degree = degree, parametric = parametric,
## : span too small.  fewer data values than degrees of freedom.

## Warning in simpleLoess(y, x, w, span, degree = degree, parametric = parametric,
## : pseudoinverse used at 0.98

## Warning in simpleLoess(y, x, w, span, degree = degree, parametric = parametric,
## : neighborhood radius 2.02

## Warning in simpleLoess(y, x, w, span, degree = degree, parametric = parametric,
## : reciprocal condition number 0

## Warning in simpleLoess(y, x, w, span, degree = degree, parametric = parametric,
## : There are other near singularities as well. 4.0804
```

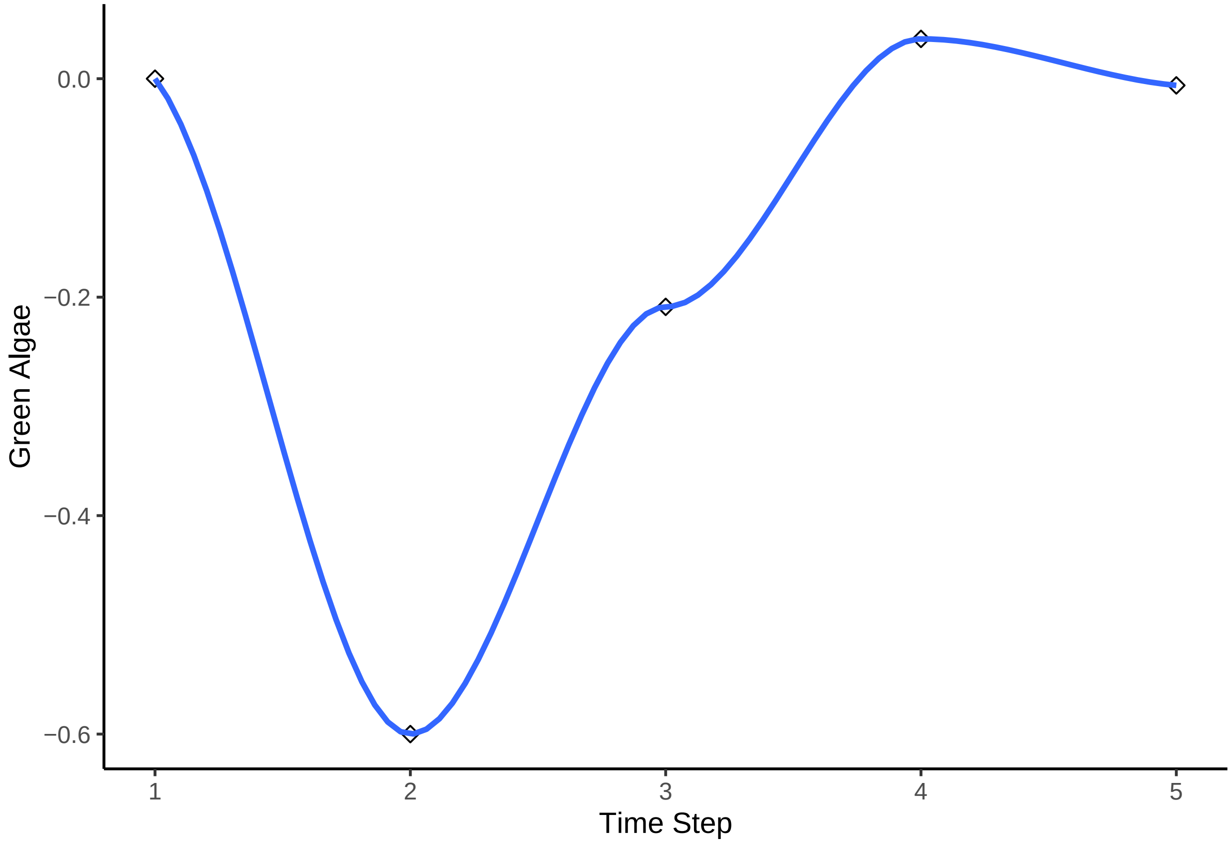


```
## ‘geom_smooth()‘ using formula = ’y ~ x’

## Warning in simpleLoess(y, x, w, span, degree = degree, parametric = parametric,
## : span too small.  fewer data values than degrees of freedom.

## Warning in simpleLoess(y, x, w, span, degree = degree, parametric = parametric,
## : pseudoinverse used at 0.98

## Warning in simpleLoess(y, x, w, span, degree = degree, parametric = parametric,
## : neighborhood radius 2.02

## Warning in simpleLoess(y, x, w, span, degree = degree, parametric = parametric,
## : reciprocal condition number 0

## Warning in simpleLoess(y, x, w, span, degree = degree, parametric = parametric,
## : There are other near singularities as well. 4.0804
```

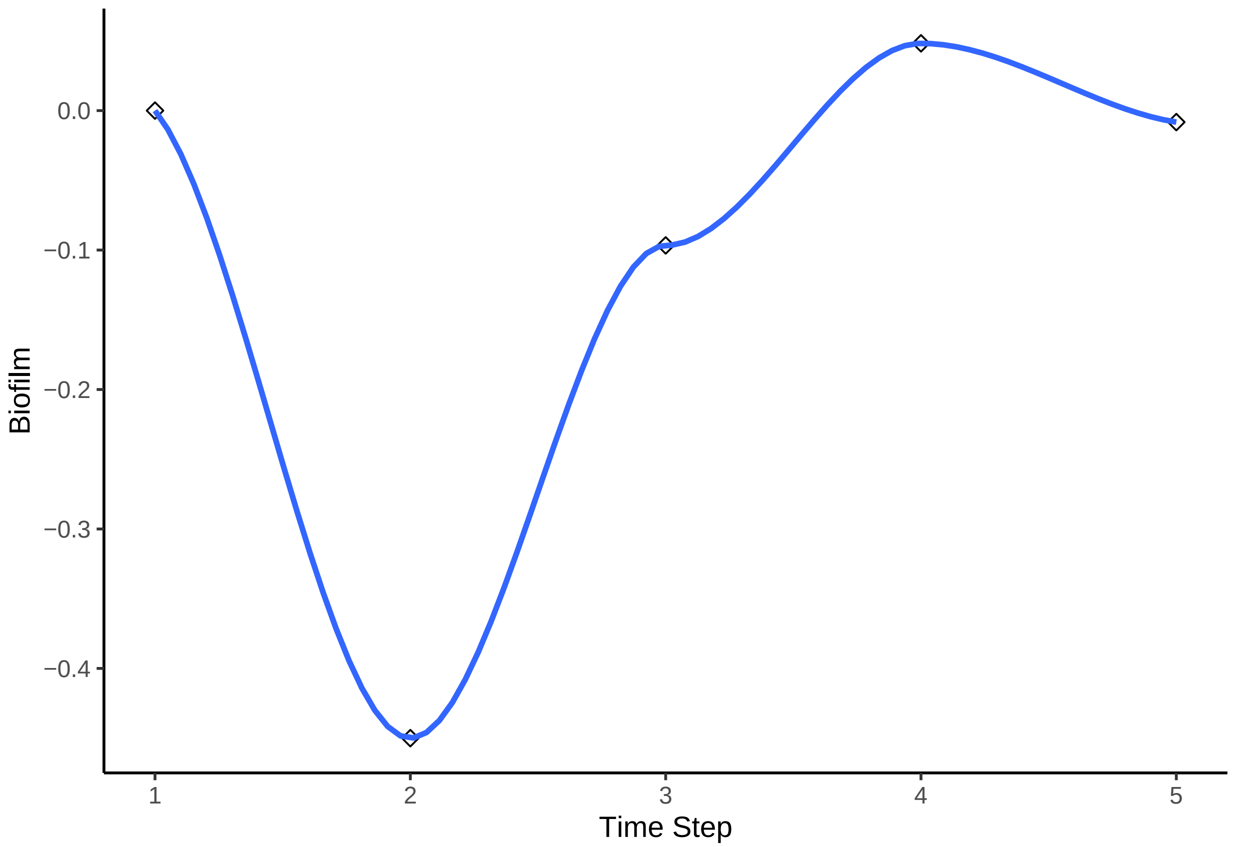


```
## `geom_smooth()` using formula = 'y ~ x'

## Warning in simpleLoess(y, x, w, span, degree = degree, parametric = parametric,
## : span too small.  fewer data values than degrees of freedom.

## Warning in simpleLoess(y, x, w, span, degree = degree, parametric = parametric,
## : pseudoinverse used at 0.98

## Warning in simpleLoess(y, x, w, span, degree = degree, parametric = parametric,
## : neighborhood radius 2.02

## Warning in simpleLoess(y, x, w, span, degree = degree, parametric = parametric,
## : reciprocal condition number 0

## Warning in simpleLoess(y, x, w, span, degree = degree, parametric = parametric,
## : There are other near singularities as well. 4.0804
```

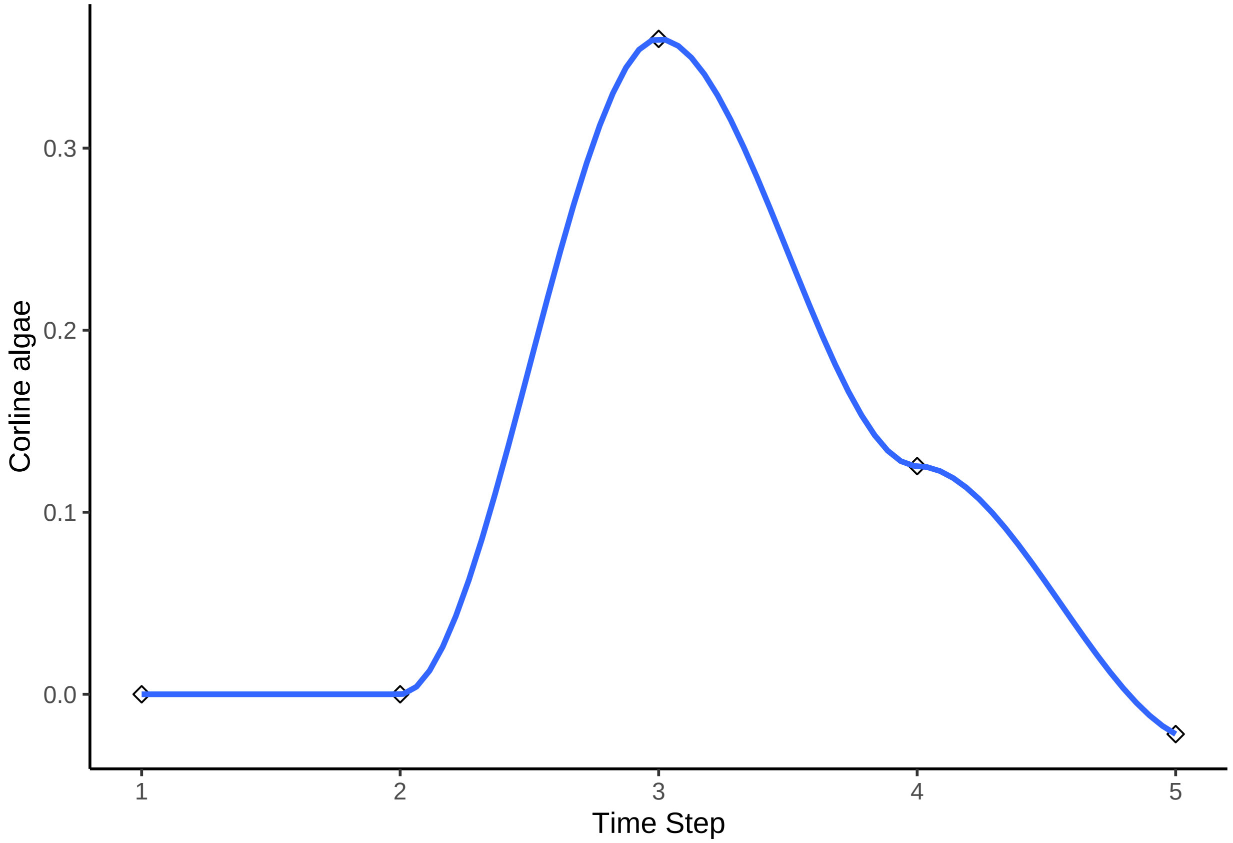


```
## `geom_smooth()` using formula = 'y ~ x'

## Warning in simpleLoess(y, x, w, span, degree = degree, parametric = parametric,
## : span too small.  fewer data values than degrees of freedom.

## Warning in simpleLoess(y, x, w, span, degree = degree, parametric = parametric,
## : pseudoinverse used at 0.98

## Warning in simpleLoess(y, x, w, span, degree = degree, parametric = parametric,
## : neighborhood radius 2.02

## Warning in simpleLoess(y, x, w, span, degree = degree, parametric = parametric,
## : reciprocal condition number 0

## Warning in simpleLoess(y, x, w, span, degree = degree, parametric = parametric,
## : There are other near singularities as well. 4.0804
```

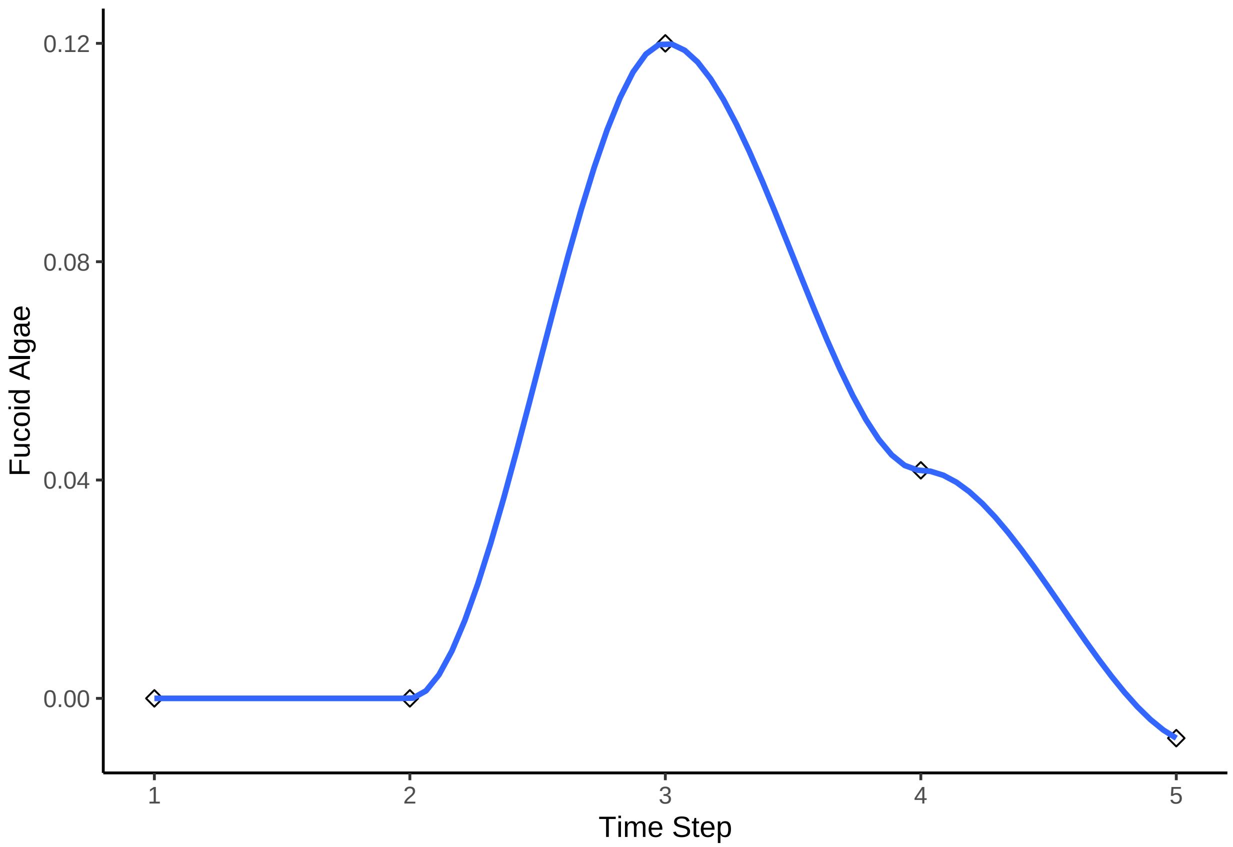


##### bbn.visualise()

This function produces a network diagram in each timestep of the model, rather than the graph produced above. The most important interactions at that timestep are listed and the colour of nodes changes from black (showing the highest increase) to white (showing the lowest increase / largest decrease)

*Required arguments*

*bbn.model* - a matrix or dataframe of interactions between different model *nodes*

*priors1* - an X by 2 array of initial changes to the system under investigation. The first column should be a -4 to 4 (including 0) integer value for each node in the network with negative values indicating a decrease and positive values representing an increase. 0 represents no change.

*Optional Arguments*

*timesteps* - default = 5. This is the number of timesteps the model performs. Note, timesteps are arbitrary and non-linear. However, something occurring in timestep 2, should occur before timestep 3.

*disturbance* - default = 1. 1 - creates a prolonged or press disturbance as per the bbn.predict() function. Essentially prior values for each manipulated node are at least maintained (if not increased through reinforcement in the model) over all timesteps. 2 - shows a brief pulse disturbance, which can be useful to visualise changes as peaks and troughs in increase and decrease of nodes can propagate through the network

*threshold* - default = 0.2. Nodes which deviate from 0 by more than this threshold value will display interactions with other nodes. As mentioned, values in these visualisation functions don't directly correspond to those in the bbn.predict() function. This value can be tweaked from 0 to 4 to create the most useful visualisations *font.size* - default = 0.7. Changes the font in the figure produced. The value here is a multiplier of the default font size used in the *igraph* package and does not correspond to the font.size argument in the bbn.timeseries() function.

*arrow.size* - default = 4. Changes the size of the arrows. Note, sizes do vary based on interaction strength, so this is a multiplier for visualisation purposes.

*Example*

```
bbn.visualise(bbn.model = my_BBN, priors1 = combined, timesteps = 5, disturbance = 2, threshold=0.05, fo
```

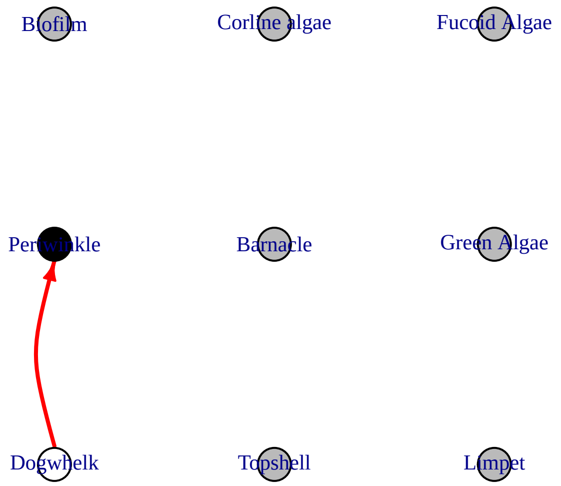


```
## NULL
```

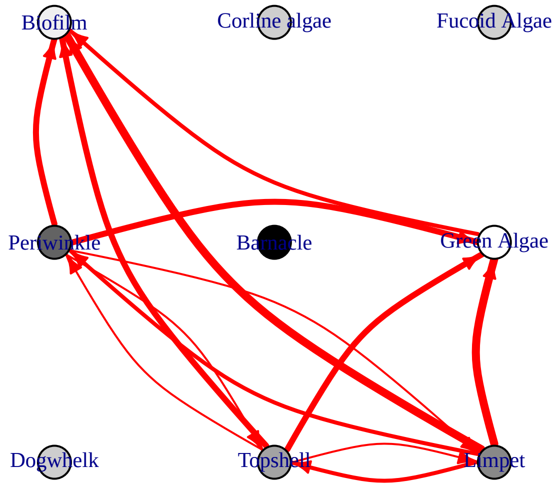
Biofilm
Corline algae
Fucoid Algae
Periwinkle
Barnacle
Green Algae
Dogwhelk
Topshell
Limpet

```
## NULL
```

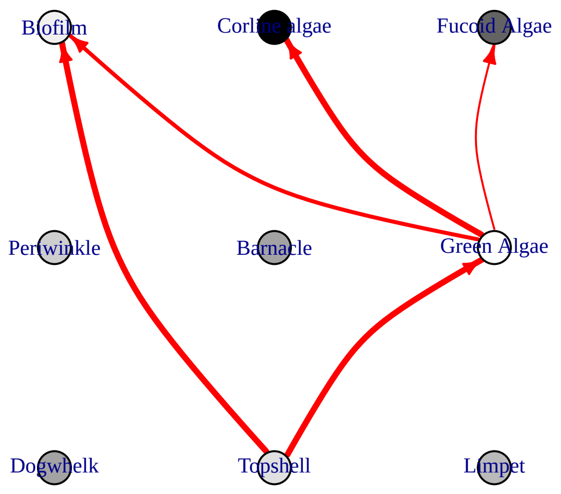

Biofilm
Corline algae
Fucoid Algae
Periwinkle
Barnacle
Green Algae
Dogwhelk
Topshell
Limpet


```
## NULL
```

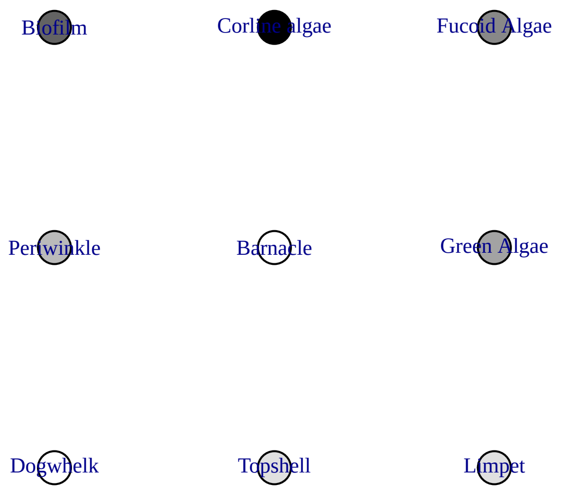
Biofilm
Corline algae
Fucoid Algae
Periwinkle
Barnacle
Green Algae
Dogwhelk
Topshell
Limpet

```
## NULL
```

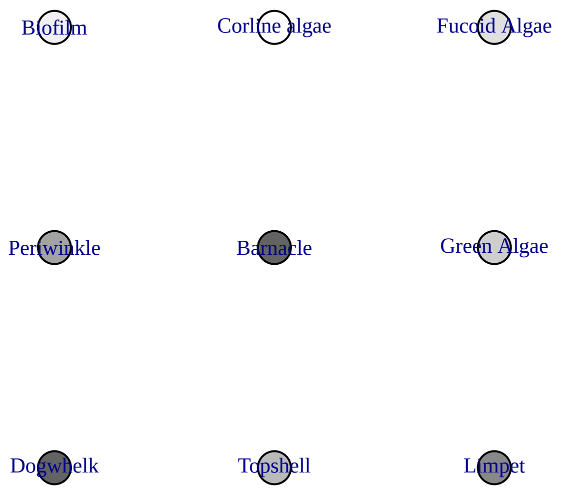


```
## NULL
```

#### Parameterisation of the model

#### bbn.sensitivity()

For some methods of model parameterisation, extensive data extraction from literature, or expert opinion can be useful. However, this is time consuming, and being aware of the most sensitive parameters in the model which may affect the desired outputs could help concentrate efforts. This function produces a list of the most important parameters / interaction strengths to examine.

*Required arguments*

*bbn.model* - a matrix or dataframe of interactions between different model *nodes*

One or more nodes (recommended no more than 3) which would be the main outcomes of interest in the model. The spelling of these nodes needs to be identical (including capital letters) to that in the imported csv file (note, you should include spaces if these are in your csv file, rather than the dot notation used once imported into R).

*Optional arguments*

*boot_max* - the number of bootstraps to perform. Suggested range for exploratory analysis 100-1000. For final analysis recommended size = 1000 - 10000 - note, this can take a long time to run. Default value is 1000.

*Example use and interpretation*

```r
bbn.sensitivity(bbn.model = my_BBN, boot_max = 100, 'Limpet', 'Green Algae')
```

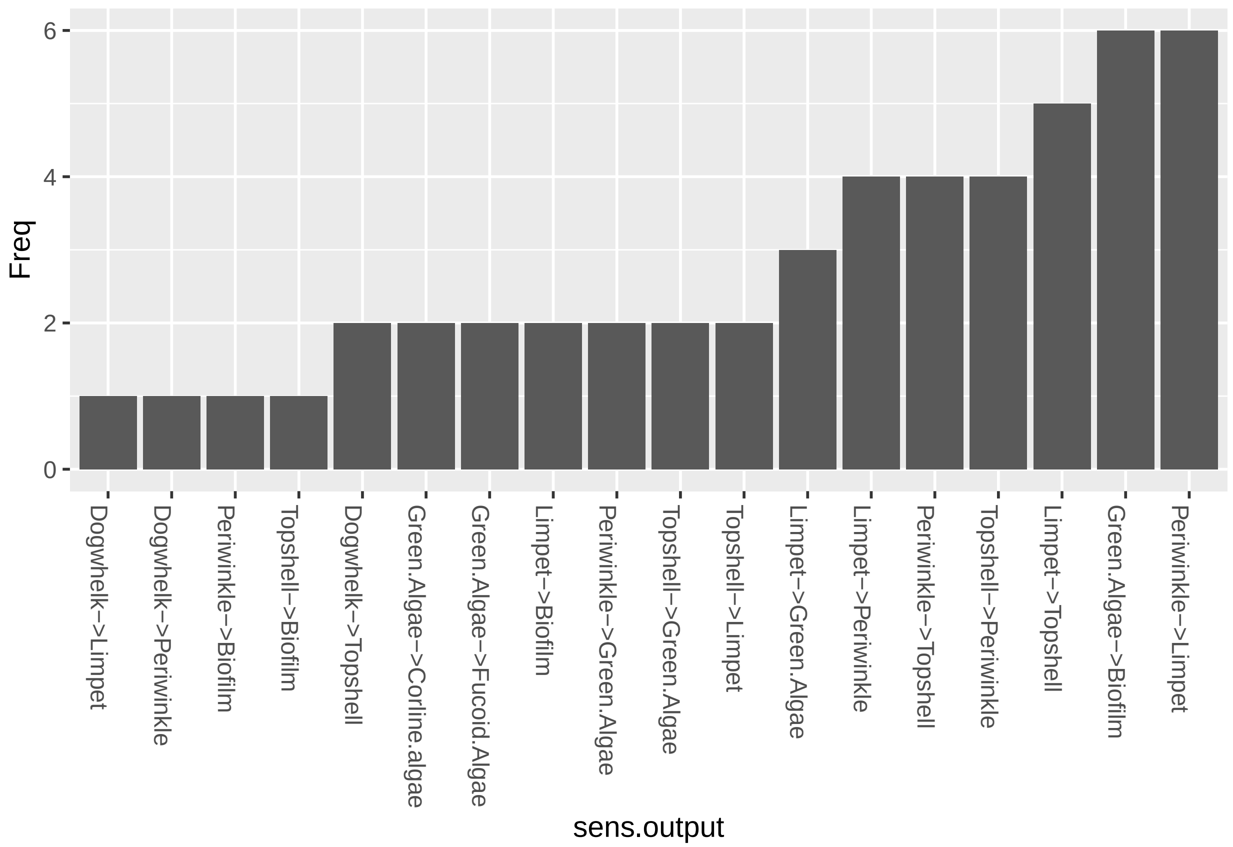


```
##                    sens.output Freq
## 1             Dogwhelk->Limpet    1
## 2         Dogwhelk->Periwinkle    1
## 3          Periwinkle->Biofilm    1
## 4            Topshell->Biofilm    1
## 5           Dogwhelk->Topshell    2
## 6   Green.Algae->Corline.algae    2
## 7    Green.Algae->Fucoid.Algae    2
## 8              Limpet->Biofilm    2
## 9      Periwinkle->Green.Algae    2
## 10       Topshell->Green.Algae    2
## 11            Topshell->Limpet    2
## 12         Limpet->Green.Algae    3
## 13          Limpet->Periwinkle    4
## 14        Periwinkle->Topshell    4
## 15        Topshell->Periwinkle    4
## 16            Limpet->Topshell    5
## 17         Green.Algae->Biofilm    6
## 18          Periwinkle->Limpet    6
```

The function works by bootstrapping with multiple changes to prior values and interaction strengths in the network. The frequency shows the number of times a modified interaction shows up as important in causing a change to the listed nodes. As such, those interactions showing as more frequent in the table or figure are likely to be most influential in any predictions. These should be subject to closer scrutiny in terms of values used. (note, this does not mean the values are incorrect or should be reduced from more extreme values - i.e. from 4 to 3, just that they should be carefully checked)

#### Visualising the network

For simple networks visualising the interactions can be useful. In some more complex cases, network diagrams will only serve to illustrate a 'complex system' exists.

#### bbn.network.diagram()

This visualises all nodes and inetactions in a network, in a similar manner to the bbn.visualise() package, other than this is the full network. Nodes can also be colour coded by theme. It is easiest to create a slightly different csv file to produce these networks, which allows for the colour coding.

```
my_network <- read.csv('RockyShoreNetworkDiagram.csv', header=T)

head(my_network)
```

```
##    id node.type   node.name Dogwhelk Topshell Limpet Periwinkle Barnacle
## 1 s01         1    Dogwhelk       NA       -1     -2         -2       -3
## 2 s02         2    Topshell       NA       NA     -1         -1       NA
## 3 s03         2      Limpet       NA       -2     NA         -2       NA
## 4 s04         2  Periwinkle       NA       -1     -1         NA       NA
## 5 s05         3    Barnacle       NA       NA     NA         NA       NA
## 6 s06         4 Green Algae       NA       NA     NA         NA       NA
##   Green.Algae Biofilm Corline.algae Fucoid.Algae
## 1          NA      NA            NA           NA
## 2          -3      -3            NA           NA
## 3          -4      -4            NA           NA
## 4          -3      -3            NA           NA
## 5          NA      NA            NA           NA
## 6          NA      -2            -3           -1
```

Note - in this file, the first column is called `id` and consists of an s and a 2 digit number relating to the node number. The second column is called `node.type` and is an integer value from 1-4. This sets the colour of the node in the network (sticking to a maximum of four colours). Here, predators, grazers, filter feeders and algae are colour coded seperately - it would be fine to change the colours, for example to ensure algae were green. The third column is the same as the first column in the standard BBN interaction csv, other than it is titled `node.name`. It is important to use these column names (including capitals and dot notation). The remainder of the columns are exactly as the standard BBN interaction csv file.

*Required arguments*

*bbn.network* - a csv file as described above, with note paid to the first three column names

*Optional arguments*

*font.size* - default = 0.7. Changes the font in the figure produced. The value here is a multiplier of the default font size used in the *igraph* package and does not correspond to the font.size argument in the bbn.timeseries() function.

*arrow.size* - default = 4. Changes the size of the arrows. Note, sizes do vary based on interaction strength, so this is a multiplier for visualisation purposes. Negative interactions are shown by red arrows, and positive interactions by black arrows *arrange* - this describes how the final diagram looks. Default is `layout_on_sphere` but `layout_on_grid` provides the same layout as in the bbn.visualise() function and ensures nodes are structured in the order specified in the network. Other layouts, including `layout_on_sphere` are more randomly determined, and better/clearer diagrams may occur if you run these multiple times. Other options are from the *igraph* package:

```
layout.sphere
layout.circle
layout.random
layout.fruchterman.reingold
```

*Examples*

```r
bbn.network.diagram(bbn.network = my_network, font.size = 0.7, arrow.size = 4, arrange = layout_on_spher
```

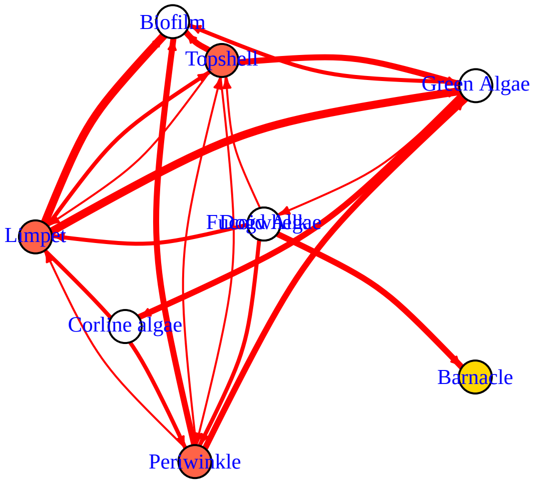


```
## NULL
```

```r
bbn.network.diagram(bbn.network = my_network, font.size = 0.7, arrow.size = 2, arrange = layout_on_grid
```

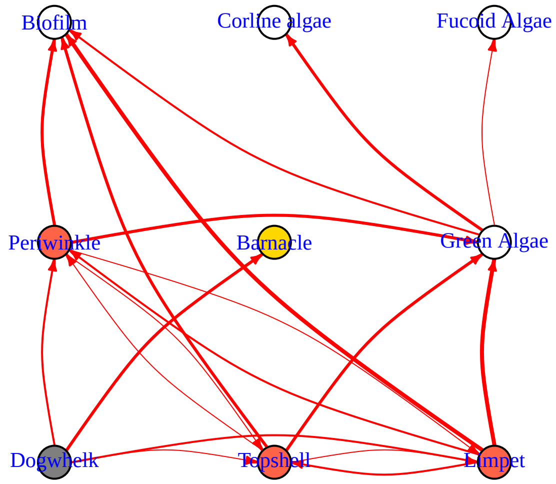


```
## NULL
```

```r
bbn.network.diagram(bbn.network = my_network, font.size = 0.7, arrow.size = 2, arrange = layout.random)
```

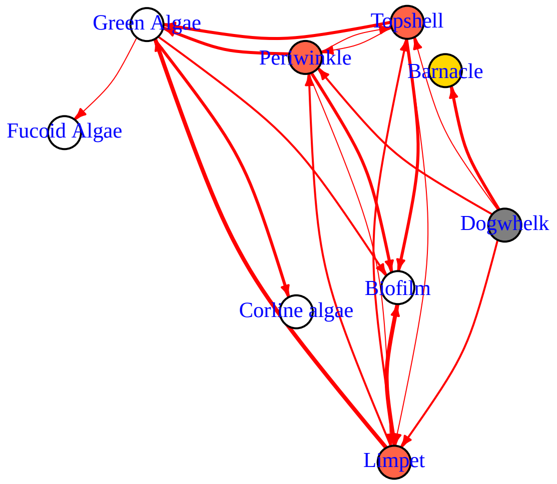


```
## NULL
```

```
bbn.network.diagram(bbn.network = my_network, font.size = 0.7, arrow.size = 2, arrange = layout.circle)
```

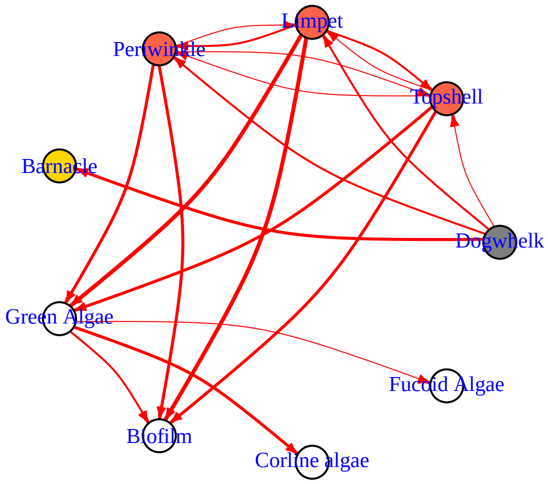
Limpet
Periwinkle
Topshell
Barnacle
Dogwhelk
Green Algae
Fucoid Algae
Biofilm
Corline algae

```
## NULL
```
